# Network organization of protein binding-site landscapes across scales

**DOI:** 10.64898/2026.10.17.759831

**Authors:** G. Giordano, A. Chernomoretz

## Abstract

Studying protein binding sites helps explain ligand recognition and selectivity, informing drug development and repurposing. Here, we combine protein language model embeddings with network analysis to investigate how local recognition environments organize across scales. We represented 121,673 non-redundant, experimentally identified binding sites using ProtT5 embeddings of ligand-contacting residues and constructed a mutual nearest-neighbor network. Embedding proximity was associated with local structural similarity, supporting the structural relevance of network relationships. Globally, the network exhibited ligand-related organization. At the mesoscopic scale, communities revealed heterogeneous landscapes within established protein classifications, with InterPro categories ranging from compact neighborhoods to dispersed recognition environments. Focusing on kinases, a major class of drug targets, we found that communities within individual proteins distinguished separate binding regions and resolved recurrent residue-defined microenvironments within broad structural classes. Local similarities also connected sites in proteins with low sequence identity and non-overlapping InterPro annotations, suggesting shared recognition features across distant proteins. Together, these results show how network organization captures both diversification within protein categories and local similarities across their boundaries. This multiscale framework complements conventional protein classifications and provides a basis for prioritizing structural and biochemical investigations of ligand cross-reactivity, selectivity, and repurposing opportunities.

## Introduction

Protein binding sites provide a local perspective on molecular recognition that complements whole-protein sequence, fold, and evolutionary classifications. Similar ligand-contacting environments can occur in different protein contexts, potentially supporting shared ligand recognition and cross-target activity. Consistent with this idea, binding-site similarity has been associated with drug promiscuity, motivating its investigation in drug repurposing, polypharmacology, and selectivity assessment (Haupt et al. 2013) . Structural comparisons have established that binding-site space contains recurrent patterns across proteins. (Kinjo and Nakamura 2009) identified approximately 3,000 structural motifs through large-scale comparison of ligand-binding sites. Network approaches extended this perspective by organizing pairwise similarities into broader relational landscapes. (Z. Zhang and Grigorov 2006) introduced binding-site similarity networks for functional classification and computational chemogenomics, while (Lee et al. 2017) identified structural communities spanning diverse ligands and protein domain contexts. These studies established a foundation for investigating molecular recognition through interconnected local environments.

Learned representations offer complementary approaches to describing these environments. Protein language models such as ProtT5 generate contextual residue embeddings that retain information useful for structural and functional prediction (Elnaggar et al., 2022). Structure-conditioned representations have also supported binding-site prediction, as demonstrated by IF-SitePred (Carbery et al., 2024), while ATOMICA learns atomic-scale representations of intermolecular interfaces across multiple molecular modalities (Fang et al., 2025). These developments motivate investigating how learned representations organize experimentally observed binding-site diversity.

Pooling sequence-derived embeddings over ligand-contacting residues provides a fixed-length representation that combines a local residue selection with broader receptor context. An important question is therefore whether relationships in this space capture recognizable binding-site features across scales. Global neighborhoods may reflect ligand-associated regularities, mesoscopic communities may reveal variation within and across protein classifications, and intra-protein subdivisions may distinguish separate binding regions or alternative contact footprints within the same pocket. Establishing these relationships is necessary to interpret embedding proximity in terms of molecular recognition.

Here, we combine ProtT5-derived binding-site embeddings with complex-network analysis to characterize experimentally resolved binding-site landscapes across scales. We construct a non-redundant atlas and a mutual nearest-neighbor network, and assess the association between embedding similarity and local structural similarity. We then examine ligand-associated neighborhood organization, recognition microenvironments within individual kinases, and the relationship between network communities and conventional sequence and InterPro classifications. Finally, we investigate local similarities between sites hosted by proteins with low sequence identity and non-overlapping InterPro annotations. This framework connects local recognition environments with broader protein-space organization and generates hypotheses for structural and pharmacological investigation.

## Materials and Methods

### Data source

We used BioLiP2 (C. Zhang et al. 2024), a curated database of biologically relevant ligand–protein interactions derived from Protein Data Bank (PDB) structures (Burley et al. 2025). The dataset was downloaded in May 2024. We retained 422,246 binding-site (BS) records from 110,159 PDB entries with a resolution of 3 Å or better. These records covered 24,334 proteins and included interactions with 42,477 distinct small molecules and 207 ions. Of the retained sites, 380,156 involved small molecules or ions, 16,440 DNA, 4,281 RNA, and 21,369 peptides.

### Binding site embeddings and network construction

Figure 1A summarizes our analysis workflow. We generated residue embeddings with the ProtT5 protein language model (Elnaggar et al. 2022). For each binding site, we used the PDB-derived receptor sequence realigned to UniProt coordinates. Gaps in the experimentally observed sequence were filled with the corresponding UniProt canonical sequence residues (see Supplementary Figure SF1). We averaged the ProtT5 residue vectors over the residues assigned to that binding site, yielding a 1,024-dimensional vector. Following this procedure we could successfully map 386,969 protein–ligand binding site sequences.

**Figure 1.**
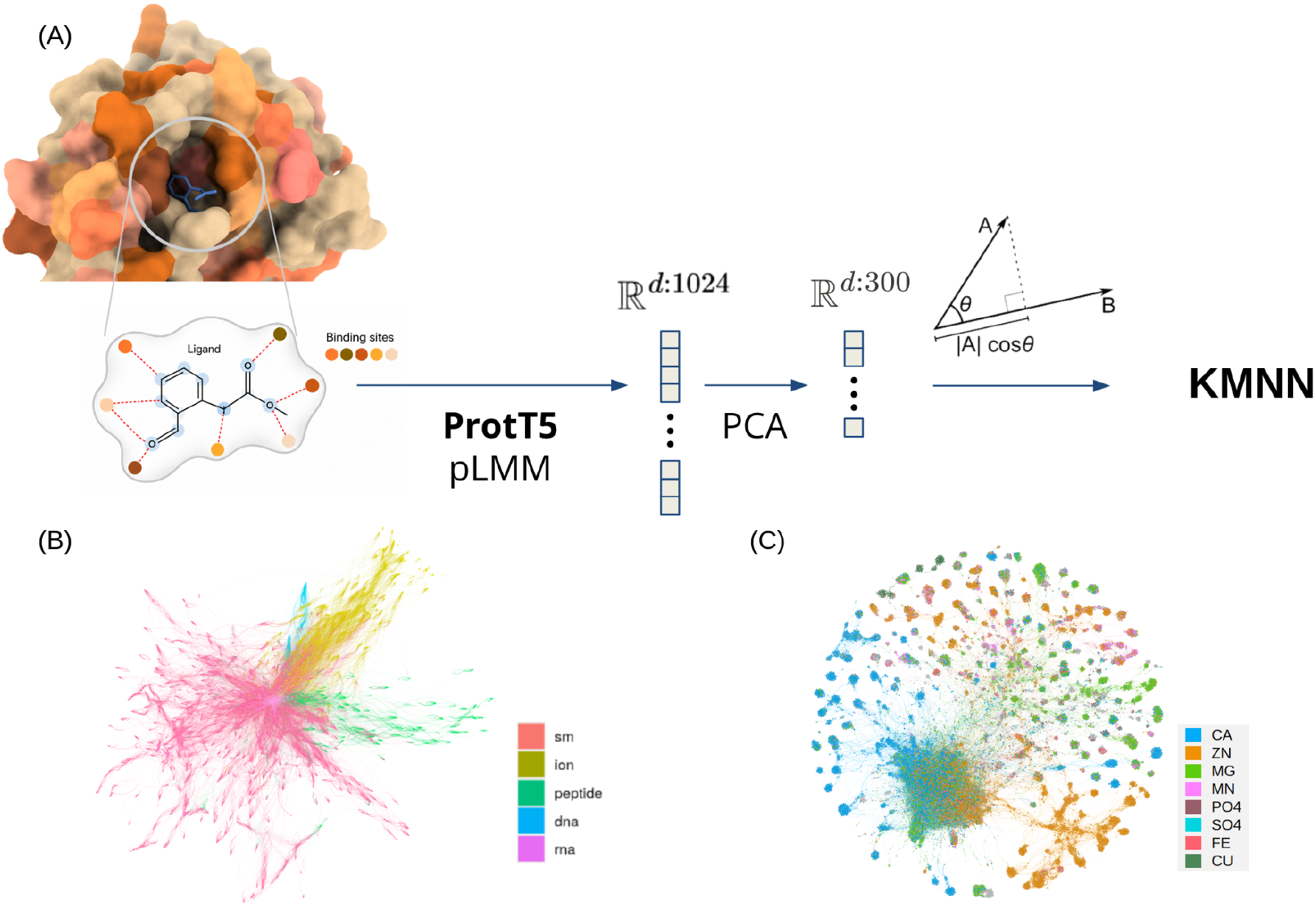
Binding-site representation and global network organization. (A) Analysis workflow. (B) Global binding-site network. (C) Ion-binding-site subgraph, with the eight most abundant ions distinguished by color.

We used mutual k-nearest-neighbor graphs to represent local relationships in BS embedding space. First, we considered principal component analysis (PCA) to reduce the embedding matrix to its first 300 components, which explained approximately 80% of the variance. Then, neighbor relationships were defined using normalized cosine similarity between the resulting vectors. Two binding sites were connected when each belonged to the other’s set of k nearest neighbors. We selected k = 30, the smallest value for which the largest connected component contained approximately 90% of the nodes. We verified that the main structural properties remained consistent at higher k values (Supplementary Figure SF2). We used this network, where each node represents an experimentally resolved protein–ligand complex, to examine the BS organization across local, mesoscopic, and global scales.

### Protein annotations and kinase binding site classification

We used InterPro version 108 annotations (Blum et al. 2025) to relate the BS landscape to protein classification. We considered homologous superfamily, family, and domain annotations as complementary descriptions of evolutionary relationships and domain architecture. We also compared network communities with protein sequence clusters defined at 30% sequence identity defined using the DIAMOND software with default parameters (Buchfink et al. 2021).

For the eight kinases with the largest numbers of binding sites—CDK2, CK2α, p38α, PIM1, EGFR, CHK1, ERK2, and AURKA—we performed a detailed, target-specific annotation in order to map their residue-contact patterns to six coarse structural classes (see Supplementary Figure SF3).

To characterize organization patterns within individual kinases, we applied Louvain community detection to the BS subgraph induced by each protein. We assessed agreement between these communities and the structural labels using purity, homogeneity, completeness, and adjusted mutual information (AMI) statistics. Purity measures the fraction of sites assigned to the dominant structural class in each community, aggregated with community-size weights. Homogeneity measures whether communities contain sites from a single structural class, whereas completeness measures whether sites from the same class are assigned to a single community. AMI summarizes agreement between the partitions after adjustment for chance. High homogeneity combined with low completeness therefore indicates that structurally coherent communities subdivide a broader structural class.

### Embedding diversity and community compaction

For each InterPro category, geometric diversity in embedding space, *G*_*div*_, was quantified as the geometric mean of *p*_*globalextent*_, the within-classification percentile of the mean pairwise embedding distances of the associated BS, and *p*_*effniches*_, the corresponding percentile of *N*eff, the effective number of Infomap communities defined as:

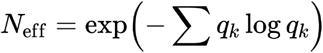

q_k_ is the fraction of binding sites assigned to the Infomap community k. N_eff_ downweights small or sparsely populated communities and represents the number of equally abundant niches required to reproduce the observed community entropy. We independently quantified the degree to which these communities accounted for global geometric variation using

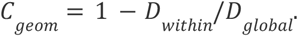

where D_global_ is the mean BS distance within a given InterPro category and D_within_ the weighted average of the mean distances of its infomap associated communities. *G*_*div*_ identifies categories that both span a broad region of embedding space and contain several substantially populated neighborhoods, whereas *C*_*geom*_ measures whether those neighborhoods constitute geometrically compact units (a.k.a. niches). Because the components of *G*_*div*_ were rank-normalized separately for domains and homologous superfamilies, its magnitude reflects relative diversity within each annotation level; *C*_*geom*_, in contrast, can be interpreted directly across levels.

## Results

### Structural calibration and a nonredundant binding site atlas

We first aimed to calibrate the distances induced by the pLMM representation by contextualizing the similarity derived in embedding space with established structural bioinformatics benchmarks. To this end, we employed ProBiS, a robust algorithm designed to detect local structural similarities (Konc et al. 2021). We uniformly sampled across embedding-distance intervals (Δ=0.05) 10000 BS pairs per bin and compared their cosine distances with local structural alignment scores. ProBiS Z-scores greater than 2 were treated as evidence of significant structural similarity (Konc et al. 2021).

As shown in Supplementary Figure SF4 the local structural similarity increased with cosine similarity in embedding space. In particular, BS pairs with cosine similarity above 0.75 showed high ProBiS similarity, whereas structural alignment scores and the proportion of successful ProBiS comparisons sharply declined below this threshold. These observations support the use of embedding neighborhoods to identify candidate relationships between structurally similar binding sites.

We then constructed a non-redundant BS library by consolidating instances of the same protein–ligand pair with cosine similarity greater than 0.9 into representative sites. This yielded 121,673 non-redundant BS vectors. The reduction of approximately 70% successfully mitigated chain redundancy and PDB multiplicity, facilitating a more streamlined and quantitative exploration of the global protein-ligand landscape.

### Local network neighborhoods show ligand and protein homophily

The mutual k-nearest-neighbor network with k = 30 contained 121,673 nodes and 1,028,709 edges (included as Supplementary Material SM1). Each node represented a non-redundant BS associated with a protein–ligand pair in a PDB entry. Ligand classes comprised small molecules, ions, peptides, DNA, and RNA, represented by 80609, 24,659, 12,596, 1,947, and 741 nodes, respectively. A further 1121 sites were associated with technical ligands, including buffers, salts, precipitants, solvents, detergents, membrane-solubilization agents, cryoprotectants, crystallization additives (Supplementary Table ST1).

The largest connected component of the graph contained 109,564 binding sites and 920,401 edges and is represented in Figure 1B. Nodes and edges were colored by ligand class, with small molecules, ions, peptides, DNA, and RNA shown in red, yellow, green, blue, and pink, respectively. It can be appreciated that the network displays a strong assortative organization with respect to ligand types, where local neighborhoods preferentially connected sites associated with the same ligand class. Ligand-class assortativity was 0.72.

Homophily was also observed within ligand classes. The ion specific BS subgraph contained 23,278 nodes and 146,146 edges (Figure 1C). Its most abundant ions included Zn (5,549 sites), Ca (5,433), Mg (4,843), and Mn (1,448). Assortativity by ion identity was 0.46, consistent with preferential connectivity between sites recognizing the same ion. Protein-identity assortativity was also positive in this network, 0.32, indicating that the host protein also contributed to local network organization.

Together, these results show that local embedding neighborhoods reflect regularities associated with ligand class, ligand identity, and host-protein identity. This homophily behavior supports an expected guilt-by-association interpretation of local connectivity, while allowing relationships between different ligand or protein classes.

### Intraprotein communities resolve kinase recognition microenvironments

We next focused on a network of small-molecule and ion-BS (comprising 94,832 nodes and 757,400 edges) to examine BS organization beyond individual neighborhoods. Within this network, we examined proteins assigned to the InterPro homologous superfamily IPR011009, the protein kinase-like domain superfamily. This group provides a relevant setting for examining recognition microenvironments because kinases regulate diverse cellular processes and are established pharmacological targets. They share a broadly related catalytic architecture but contain multiple ligand-binding regions and conformational states. The structural classes used in this analysis are described in Methods and Supplementary Figure SF3.

The kinase subgraph contained 5,784 small-molecule binding sites from proteins assigned to IPR011009 (Figure 2). Colors indicate intra-protein Louvain communities for the eight kinases with the largest numbers of represented sites: CDK2 (369), CK2α (253), p38α (206), PIM1 (169), CHK1 (150), EGFR (150), ERK2 (146), and AURKA (143).

**Figure 2.**
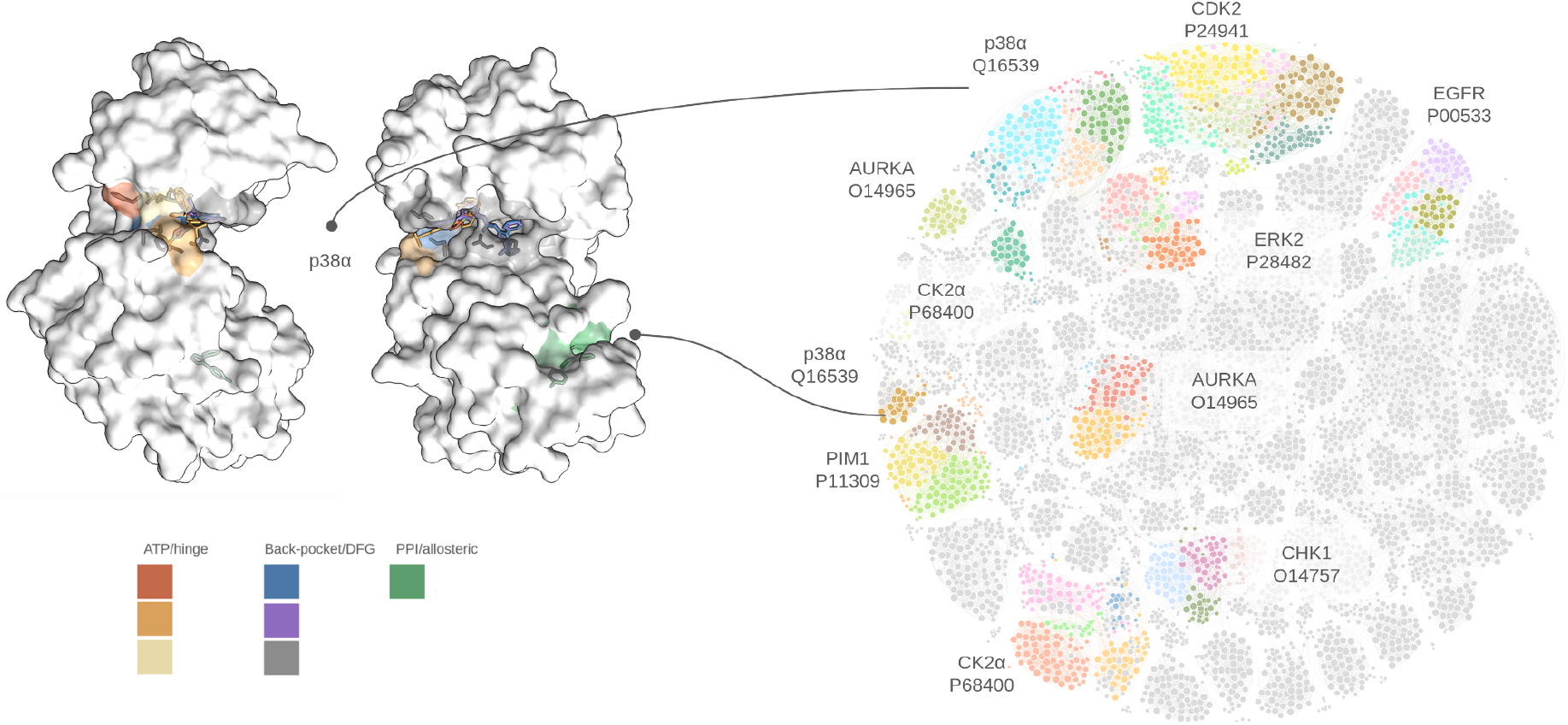
Binding-site community organization in kinases. The eight kinases with the largest numbers of binding sites are highlighted in the kinase mutual k-nearest-neighbor network (k = 30). Colors identify intra-protein Louvain communities. Inset: Binding-site patches associated with different intra-protein Louvain communities in p38α are shown with different colors.

We compared the Louvain communities of these eight kinases with their independently assigned structural classes, getting weighted purity and homogeneity values of 0.984 and 0.828 respectively. Observed purity exceeded the within-protein label-permutation expectation of 0.812 ± 0.004 (p < 0.001). Completeness was substantially lower (0.262), indicating that sites belonging to the same structural class, particularly ATP/hinge, remained distributed across multiple Louvain communities. This pattern is consistent with a hierarchy in which network communities resolve finer recognition environments within coarse structural classes (see Supplementary Figure SF5).

Figure 2 inset shows the specific case of p38α. where communities displayed a strong correspondence with ATP/hinge, back-pocket/DFG-associated, and PPI/allosteric recognition regions, including the lipid-binding pocket. In this case homogeneity was 0.969 and AMI was 0.720, indicating substantial agreement between the community and structural partitions.

Supplementary Table ST2 reports purity, homogeneity, completeness, and AMI for each kinase. Together, these measures distinguish correspondence with coarse structural classes from subdivision within a class. The eight kinases showed different patterns of agreement with coarse structural classes. CK2α, p38α, and AURKA showed clear separation of canonical ATP-site and noncanonical recognition regions, including back-pocket/DFG-associated, αD, or PPI/allosteric sites, depending on the kinase. ERK2 and CDK2 showed substantial subdivision within the canonical ATP-binding landscape; EGFR also showed stronger structural homogeneity when restricted to high-confidence annotations. CDK2 was particularly informative: one community was associated with the αC-proximal class, whereas six communities consisted exclusively of high-confidence ATP/hinge sites, with an effective number of approximately 5.7 ATP/hinge communities. CHK1 and PIM1 also contained several communities within the ATP/hinge class, but their lower agreement scores indicate that the coarse labels explain relatively little of these subdivisions (see Sup Table ST2).

Thus, intra-protein community detection recovered both separation between distinct binding regions and recurrent subdivision within canonical ATP-binding sites. The latter is consistent with differences in residue-contact footprints or subpocket usage; although contribution from conformational states would require coordinate-level assessment.

### Binding-site embeddings reveal common diversification regimes across InterPro annotation levels

We next examined how BS organization in embedding space relates to established evolutionary, domain-level, and structural classifications, and whether it reveals additional relationships that may reflect shared features of local pocket chemistry. To this end we relied on the infomap algorithm (Rosvall and Bergstrom 2008) to get a proxy of the embedding similarity landscape of the sm-ion network at the meso-scale level. The draft analysis reported 4,243 communities, comprising 799 singletons, 932 communities with 2–9 nodes, 2,023 with 10–49 nodes, and 489 with 50–327 nodes. We compared these communities with four complementary classifications: InterPro homologous superfamilies, domains, and families, and DIAMOND sequence clusters defined at 30% protein sequence identity.

We observed that Infomap communities were compact in embedding space across the entire community-size range (Supplementary Figure SF6A). In particular, the median within-community diversity values remained well below the median within-class InterPro category diversity for every size bin. This pattern suggested that conventional protein classes could span multiple embedding neighborhoods. Direct comparisons confirmed that the mean within-community diversity was systematically lower than category-level values in 95–98% of multi-community categories across the four classification schemes (Supplementary FigureSF6 3B).

We then examined how individual homologous superfamilies occupied the embedding space. Figure 3A illustrates two possible patterns. A broadly dispersed superfamily can be divided into several compact communities, whereas a superfamily with more restricted dispersion may show little further compaction within communities. The number of communities alone therefore does not describe how clearly separated their embedding neighborhoods are. In Figures 4B and 4C we characterized these patterns for domain and superfamily categories using the geometric compaction index C_geom_, the embedding diversity index G_div_, the rank-normalized effective community number, p_eff_niches_ and similarity diversity p_sim_spread_ defined in Methods.

**Figure 3.**
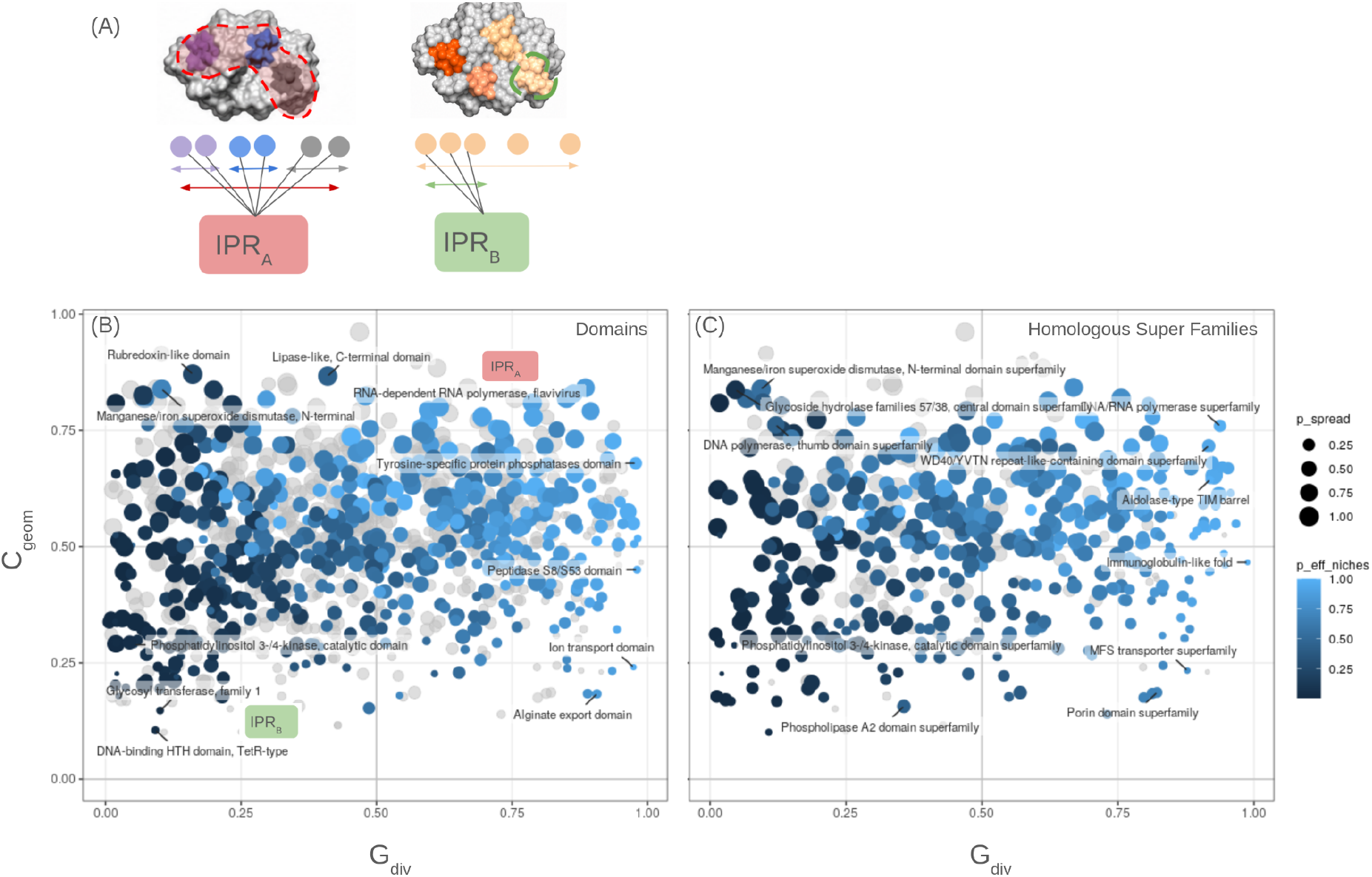
Binding-site geometric diversity and niche compaction across InterPro annotation levels. (A) Schematics illustrate the conceptual decomposition of an InterPro category into distinct local binding-site environments. (B) InterPro domains were positioned according to their geometric diversity index, and geometric compaction. Point color represents the percentile of the effective number of niches and point size the percentile of pairwise-similarity heterogeneity. Grey points denote lower-confidence categories with fewer than 10 binding sites per effective niche. Representative annotations highlight four recurrent regimes: broad diversity organized into compact niches, broad but diffuse diversity, restricted but discretely structured variation, and restricted weakly structured landscapes. (C) Homologous superfamily distribution in the G_div_ vs C_geom_ plane.

Joint analysis of InterPro domains and homologous superfamilies revealed four recurrent modes of BS organization in embedding space (Fig. 3B and C). As stated in Material and Methods, geometric diversity, *G*_*div*_, quantifies the combination of global embedding-space extent and effective niche multiplicity, whereas *C*_*geom*_ measures the reduction in pairwise distance obtained after conditioning on Infomap community membership. Importantly, the same qualitative regimes were observed at both annotation levels, indicating that the organization detected by the embeddings is not restricted to broad evolutionary classifications.

Categories with high geometric diversity and high compaction contained multiple well-separated and internally coherent BS states. This regime included the DNA/RNA polymerase homologous superfamily and the tyrosine-specific protein phosphatase domain, consistent with discrete specialization of local recognition environments around conserved catalytic frameworks. In contrast, high-diversity but weakly compacted categories, exemplified by the MFS transporter superfamily and the peptidase S8/S53 domain, occupied broad regions of embedding space without resolving into sharply isolated communities, suggesting more continuous diversification of substrate-recognition environments or conformational states.

The opposite pattern was observed for categories with low global diversity but strong compaction. The Mn/Fe superoxide dismutase superfamily and the DNA-polymerase thumb domain occupied relatively restricted regions of embedding space but contained a small number of well-defined local states, consistent with strong physicochemical or structural constraints permitting only limited discrete variation. Finally, the phospholipase A2 superfamily and PI3/4-kinase catalytic domain showed both low geometric diversity and weak compaction, indicating comparatively conserved local binding environments with little additional structure recovered by Infomap partitioning.

Similar patterns emerged for domains and homologous superfamilies, suggesting that annotation breadth alone does not explain the observed BS organization. Superfamilies encompassed both distinct embedding niches and more continuous variation, while individual domains could also contain considerable BS heterogeneity. InterPro annotations therefore provide a structural and evolutionary context, complemented by the embedding landscape’s finer description of variation in local ligand-recognition environments.

### Local binding site similarity across globally distant proteins

We next tested whether embedding similarity connected binding sites in proteins with low global similarity. In this analysis we retained edges between small-molecule binding sites, required low pairwise sequence similarity (identity < 0.3 and min_overlap>0.5) and non-overlapping InterPro annotations for the host proteins. The filtered network contained 1,823 nodes and 1,904 edges, distributed across 378 connected components (Supplementary Figure SF7 A). Selected components illustrated different potential sources of local similarity, including common cofactor-recognition features and similarities between chemically heterogeneous binding environments (Supplementary Figure SF7 B). Below, we briefly describe three of them to illustrate how these structures could suggest potentially transferable local recognition features.

Component 288 was an example of cross-family similarity. It contained binding sites from the class B1 GPCR GLP1R and four class A receptors: CB1, NTSR1, SUCNR1, and GPR174. These receptors share a seven-transmembrane core architecture but belong to distinct GPCR classes with different extracellular organization. This component therefore illustrates BS embedding similarity across sequence-divergent receptors within a shared structural framework. Noticeably, GLP1R was represented by several structurally characterized agonists, whereas the class A receptors were represented by chemically and pharmacologically diverse ligands. . Thus, the component is unlikely to reflect a shared ligand identity and instead suggests partial conservation or convergence of local transmembrane recognition environments Component 11 provided another heterogeneous example. It connected binding sites from enzymes in different metabolic pathways, including mammalian aldolase and phosphoglucomutase, bacterial phosphofructokinase, cofactor-independent phosphoglycerate mutase, and enzymes involved in organic-acid metabolism. The ligands included phosphorylated sugars, phospho-acids, and other polar metabolites or inhibitors. No single ligand accounted for the component. Recognition of polar or charged metabolites offers a possible explanation for the shared embedding features, but physicochemical convergence remains an interpretation requiring coordinate-level validation .

A related but pharmacologically more diverse pattern was observed in component 53. It connected phosphoacetylglucosamine mutase AGM1 from Candida albicans, ATIC enzymes involved in de novo purine biosynthesis, and ornithine decarboxylase from Trypanosoma brucei. The proteins belong to different structural and functional families, yet their ligand-contacting environments were connected in the network. Shared recognition features for polar or charged compounds offer a possible explanation. This component also illustrates the limits of pharmacological extrapolation: eflornithine inhibition depends on the catalytic machinery of ornithine decarboxylase, and BS embedding similarity does not establish per-se inhibition of the other proteins.

These examples suggest that local BS similarity across globally distant proteins can arise from different biological relationships (see Sup Map for details). By connecting sites with similar embeddings under stringent protein-level filters, the network provides a basis for investigating shared recognition features and their possible implications for off-target interactions and selectivity. Further structural and biochemical analyses are needed to determine whether these hypotheses based on the BS embedding space reflect recognition of common chemical motifs, remote evolutionary relationships, or physicochemical convergence, and whether they translate into pharmacologically relevant cross-reactivity.

## Discussion

Our results show that protein language model embeddings, combined with similarity networks, provide a multiscale representation of experimentally observed BS landscapes. Local neighborhoods exhibited ligand-related homophily, intra-protein communities distinguished recognition environments, for instance within individual kinases, and mesoscopic partitions described BS variation within and across conventional protein classifications. Similarity relationships also persisted between sites hosted by proteins with low sequence identity and non-overlapping InterPro annotations. Together, these findings support a view of molecular recognition in which protein-level organization and local BS variation contribute at different scales.

The representation combines experimentally defined ligand-contacting residues with sequence-derived contextual embeddings. Although three-dimensional coordinates determine which residues constitute a binding site, the resulting vectors do not explicitly encode their spatial arrangement or the chemical structure of the bound ligand. The observed association between embedding similarity and ProBiS structural similarity therefore provides an important empirical basis for interpreting this representation. Likewise, ligand-class homophily indicates that the pooled residue embeddings retain information associated with different recognition environments, even though ligand identity is not directly included in the vectors. These observations support the use of embedding neighborhoods to prioritize candidate BS relationships, while leaving their detailed structural interpretation to subsequent analysis.

The kinase analysis illustrates how network organization can resolve variation at a finer scale than protein identity or broad pocket classification. Intra-protein communities separated distinct recognition regions in proteins such as p38α, CK2α, and AURKA, while also repeatedly subdividing the canonical ATP/hinge class. The combination of high structural homogeneity and lower completeness is consistent with recurrent residue-defined recognition microenvironments nested within broader binding regions. This interpretation accommodates both anatomically distinct pockets and alternative contact footprints within the same catalytic cleft, without requiring a one-to-one correspondence between communities and physical cavities. The origin of these finer subdivisions nevertheless requires careful consideration. Different ligand-contacting residue sets can reflect alternative subpocket usage, ligand extent, or changes in pocket accessibility. Conformational differences may contribute indirectly when they alter the residues contacting a ligand, but the representation does not directly resolve conformational states: for an identical receptor sequence and an identical set of selected residues, the pooled embedding remains unchanged regardless of their three-dimensional arrangement. Consequently, subdivision of ATP/hinge sites is evidence of heterogeneity in their sequence-contextualized contact footprints, rather than direct evidence of distinct conformational states. Coordinate-level comparisons will be needed to determine which communities correspond to reproducible structural or mechanistic differences.

At the mesoscopic scale, the joint analysis of geometric diversity and community compaction provides a useful description of how BS variation is distributed within protein categories. A category can span a broad region of embedding space while resolving into compact neighborhoods, or exhibit similarly broad variation with weaker internal separation. Conversely, restricted global variation can coexist with discrete local subdivisions. The occurrence of these patterns at both domain and homologous-superfamily levels indicates that annotation breadth alone does not account for the observed organization. InterPro annotations and embedding neighborhoods thus provide complementary descriptions: the former establish evolutionary and domain context, whereas the latter describe variation among the represented ligand-contacting environments.These geometric patterns should, however, be interpreted as descriptive properties of the representation. Categories containing compact neighborhoods offer testable hypotheses about localized recognition similarities, but whether community membership predicts shared ligand activity more effectively than protein classifications remains to be established.

The filtered network extends this perspective to BS in globally distant proteins. Its components suggest several possible explanations for local similarity, including recognition of related chemical groups, remote evolutionary relationships, and physicochemical convergence. These explanations need not be mutually exclusive. Low sequence identity and non-overlapping InterPro annotations define an operationally distant set of protein pairs; they do not demonstrate the absence of homology. Similarly, the GPCR component illustrates relationships across divergent receptor classes within a shared architectural framework, rather than establishing similarity between evolutionarily unrelated proteins. The metabolic-enzyme components raise complementary hypotheses about recognition features associated with polar or charged compounds.

For drug discovery, these relationships provide a basis for prioritizing investigations of ligand transfer, off-target binding, and selectivity. A connection between two binding sites may motivate testing whether compounds recognizing one environment can engage the other, or whether shared subpocket features create a potential selectivity challenge. Such transfer cannot be inferred from embedding similarity alone. Similarity may involve only part of the recognition environment, and productive binding or inhibition can depend on additional geometric, chemical, and mechanistic requirements. Moreover, connected components summarize chains of local relationships; they do not imply that every pair of constituent sites is similar or pharmacologically interchangeable.

Several limitations define the scope of the atlas. Its coverage reflects the proteins, ligands, and binding modes represented in experimentally determined complexes. Redundancy reduction mitigates repeated observations but does not remove unequal sampling across targets or chemical series. The resulting landscape therefore describes observed binding sites rather than the full repertoire of possible recognition environments. Contextual residue embeddings may also retain substantial host-protein information, so proximity can reflect a combination of local recognition features and broader sequence context. In addition, mutual-nearest-neighbor sparsification makes connectivity dependent on local sampling density: the absence of an edge should not be interpreted as evidence of molecular incompatibility. Although the main topological properties were stable at higher neighborhood sizes, individual community boundaries and candidate relationships may remain sensitive to representation and graph-construction choices.

Future validation should therefore connect network-derived hypotheses to independent structural and pharmacological measurements. Comparisons of pocket geometry, interaction patterns, ligand chemistry, and measured cross-target activity could establish which embedding relationships are most informative for molecular recognition. Evaluation on held-out proteins and chemical series would further clarify whether these relationships generalize beyond the sampling structure of the atlas. Such analyses would also help determine when protein-context information is useful and when it obscures transferable local features. The present framework provides an organized starting point for these investigations, linking broad protein-space organization to experimentally grounded local recognition environments.

## Conclusions

We developed a network representation of experimentally resolved binding sites using contextual residue embeddings from ProtT5. The resulting landscape captures ligand-associated organization, resolves recognition microenvironments within individual proteins, and describes BS heterogeneity that is not fully accounted for by conventional protein classifications. It also identifies local similarity relationships across proteins selected for low sequence identity and non-overlapping annotations. These findings establish a framework for organizing and prioritizing hypotheses about molecular recognition across scales. Its value for drug repurposing, off-target assessment, and selectivity analysis will depend on determining which network relationships translate into independently validated structural and pharmacological similarities.

## Acknowledgments

This work was supported by Consejo Nacional de Investigaciones Científicas y Técnicas (fellowship to GG and position to AC), University of Buenos Aires via UBACyT grant 2022-20020220100181BA and Fundación Instituto Leloir.

