## supplementary figures an tables for "Network organization of protein binding-site landscapes across scales"

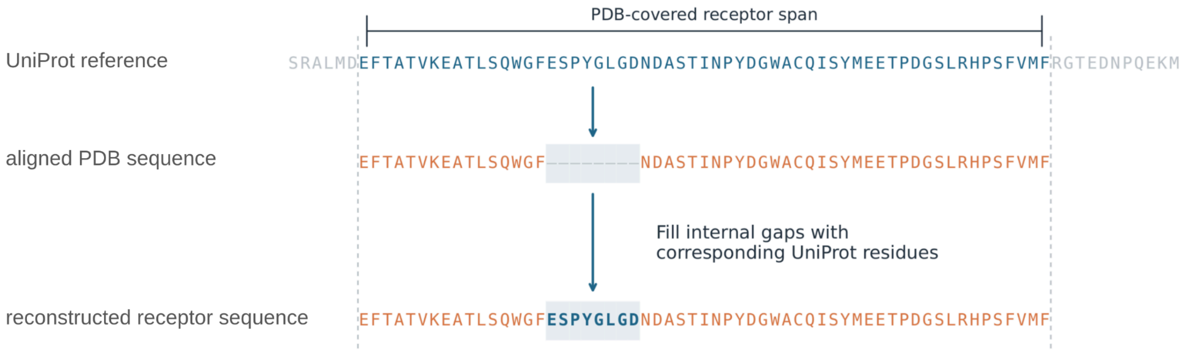


**Fig SF1:** Receptor sequence reconstruction for ProtT5. The PDB sequence is first aligned to UniProt, then internal gaps are filled within the receptor span.

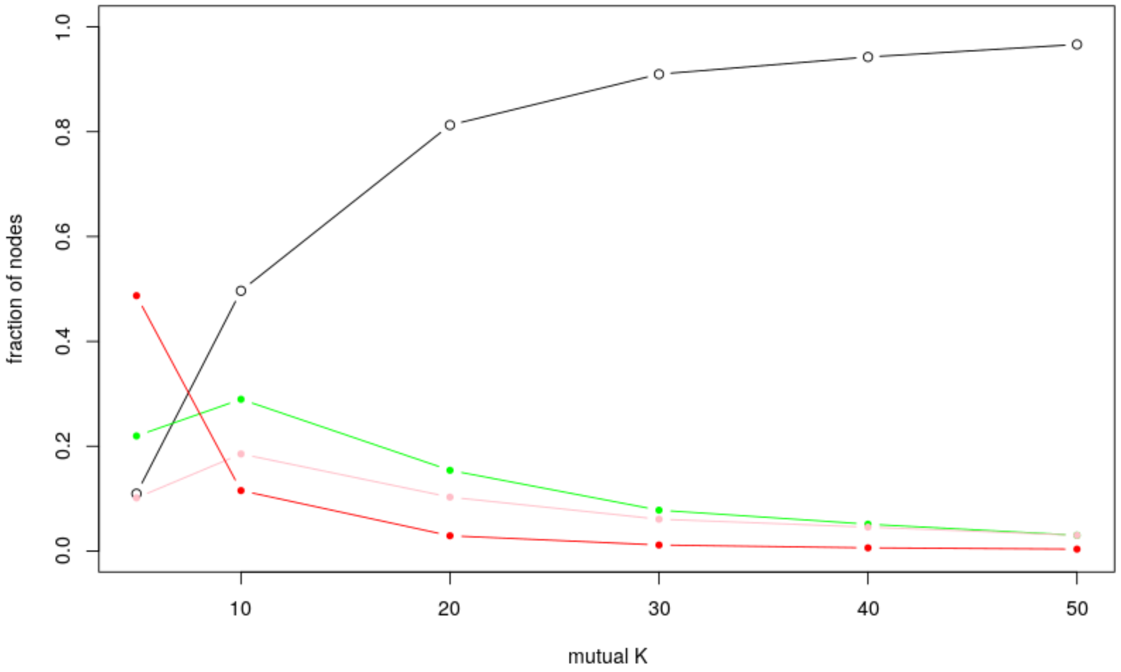


Fig SF2 Fraction of nodes in the giant component, and fraction of nodes in intermediate mass components, larger than 50 and 20 vertices, as a fusion of the parameter mutual-K, are displayed as black, pink and green lines respectively. The red line depicts the fraction of nodes in light components (mass lower than 10 nodes) as a function of the mutual-K parameter.

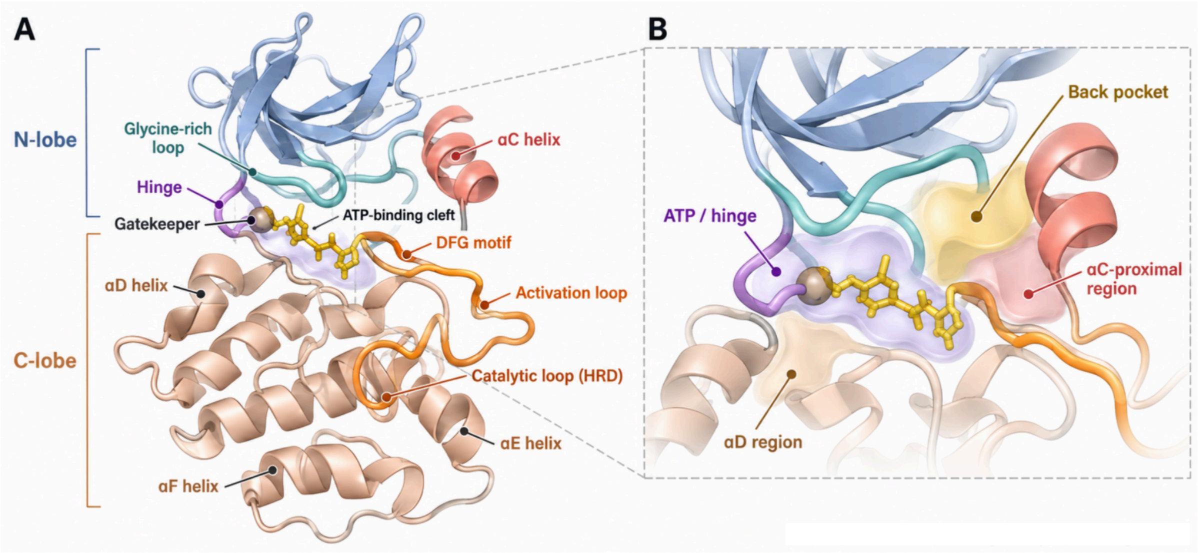


**Fig SF3** (A) Schematic representation of the conserved bilobal architecture of eukaryotic protein kinases, comprising a β-sheet-rich N-terminal lobe and a predominantly α-helical C-terminal lobe. The ATP-binding cleft lies between the two lobes, adjacent to the hinge, glycine-rich loop, gatekeeper residue, and αC helix. Additional structural elements include the DFG motif, activation loop, HRD-containing catalytic loop, and αD–αF helices. (B) Enlarged conceptual view highlighting the **ATP/hinge region**, **back pocket**, and regions **proximal to αC** and **αD**. The illustration summarizes relative spatial relationships rather than an experimentally determined structure; pocket geometry and accessibility vary among kinases and conformational states.

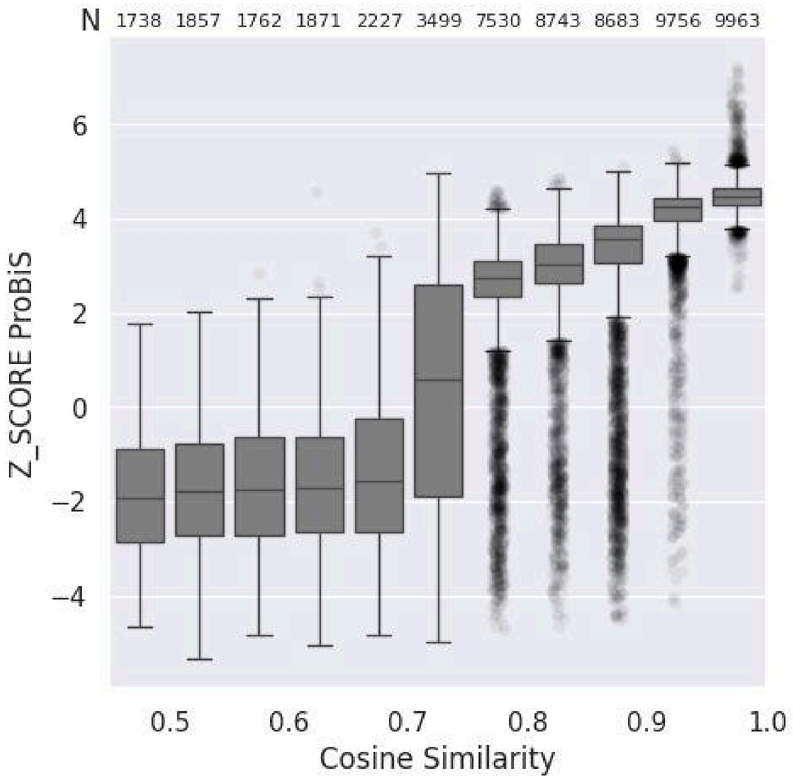


**Fig SF4**: Distribution of ProBiS alignment scores for binding-site pairs at varying embedding space distances. Each bin represents up to 10,000 sampled pairs with a cosine distance step of 0.05; the count of successful alignment estimations is indicated at the top.

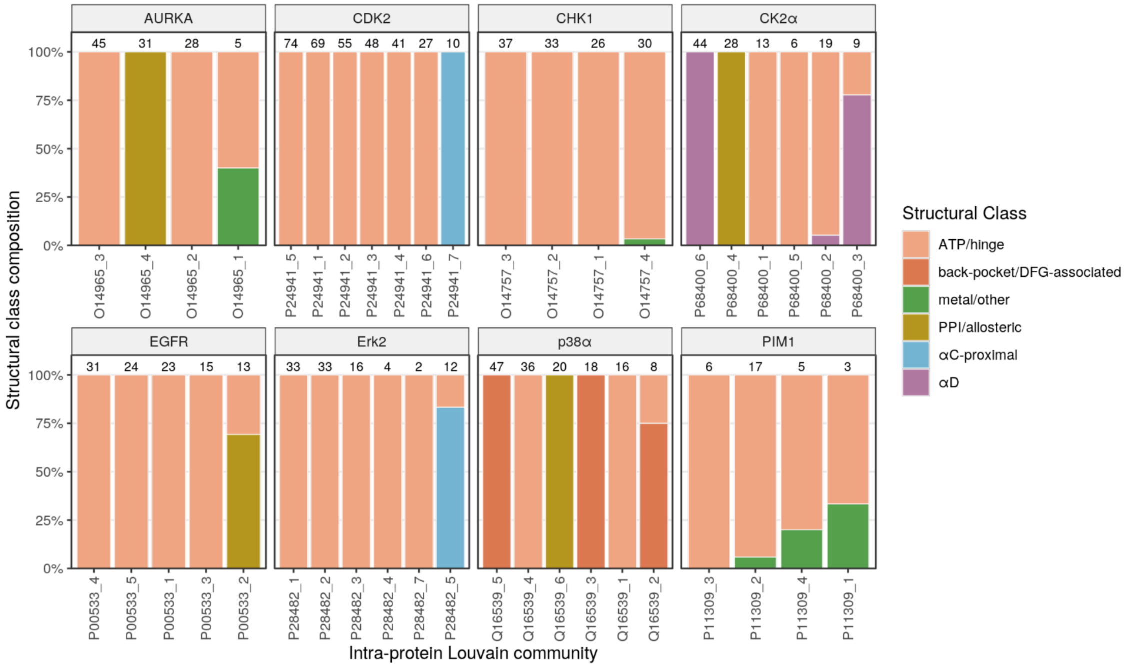


Fig SF5 Structural composition of intra-protein Louvain communities for the eight kinases with the largest numbers of binding sites. Each bar represents a community and shows the fraction of binding sites assigned to each structural class. Numbers above bars indicate the number of represented sites. The community compositions show a predominance of structurally homogeneous groups. At the same time, the ATP/hinge class was repeatedly distributed across several communities within a protein.

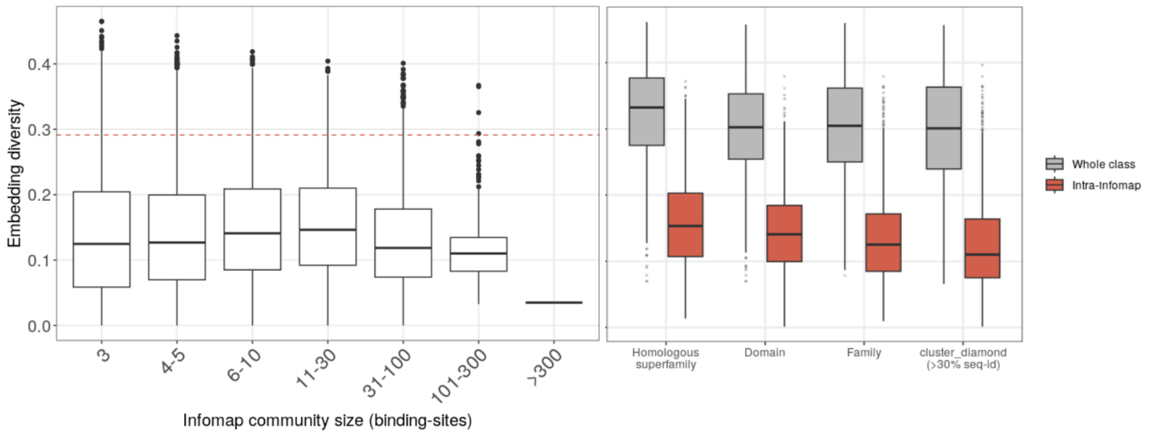


Fig SF6. Embedding diversity of Infomap communities and protein classes. (A) Distribution of within-infomap community embedding diversity values, grouped by community sizes. The horizontal red dashed line marks the reference median within-InterPro class dispersion. (B) Category-level diversity and mean within-Infomap-community diversity for InterPro homologous superfamilies, domains, families, and DIAMOND sequence clusters. Communities were more compact than their parent classes in 95–98% of multi-community categories.

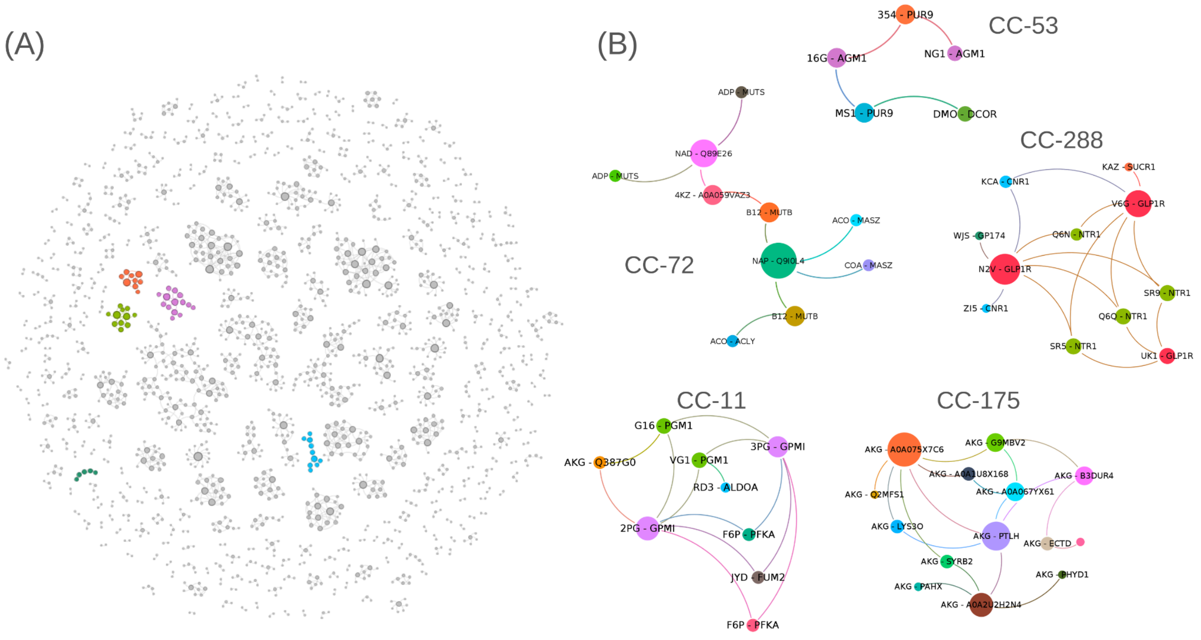


Fig SF7 (A) Connected components of the binding-site network restricted to globally distant host proteins.Components 72, 53, 288, 11, and 17, described in the text, are colored (B) Panels show in detail the selected components. Nodes are colored by UniProt identifier, and node size is proportional to degree. Labels show the ligand Chemical Component Dictionary (CCD) code and the UniProt protein name.

| CCD | Name |
|---|---|
| ALY | N6-Acetyl-L-lysine |
| C5P | Cytidine 5-monophosphate |
| GSH | Glutathione |
| SER | L-Serine |
| TMP | Thymidine 5-monophosphate (dTMP) |
| TPO | O-Phospho-L-threonine |
| TTP | Thymidine 5-triphosphate (dTTP) |
| TYS | O-Sulfo-L-tyrosine |
| EEE | Ethyl acetate |
| ARF | Formamide |
| CCN | Acetonitrile |
| DMS | Dimethyl sulfoxide (DMSO) |
| EOH | Ethanol |
| GOL | Glycerol |
| MOH | Methanol |

Sup **Table ST1. Technical ligands**

| Protein | Num BS | Num Louvain | Num struc labels | Purity | Homogeneity | Completeness | AMI |
|---|---|---|---|---|---|---|---|
| CK2α P68400 | 211 (119) | 6 | 4 (3) | 0.976 (0.975) | 0.918 (0.932) | 0.548 (0.626) | 0.678 (0.740) |
| p38α Q16539 | 198 (145) | 6 | 4 (3) | 0.955 (0.986) | 0.874 (0.969) | 0.503 (0.583) | 0.629 (0.720) |
| AURKA O14965 | 129 (109) | 4 | 5 (3) | 0.868 (0.982) | 0.714 (0.955) | 0.563 (0.537) | 0.612 (0.678) |
| ERK2 P28482 | 117 (100) | 7 | 2 (2) | 0.983 (0.980) | 0.842 (0.834) | **0.148 (0.182)** | 0.229 (0.278) |
| CDK2 P24941 | 343 (324) | 7 | 3 (2) | 0.985 (1.000) | 0.793 (1.000) | **0.078 (0.075)** | 0.126 (0.131) |
| EGFR P00533 | 145 (106) | 5 | 4 (2) | 0.759 (0.962) | **0.313** (0.739) | **0.153 (0.138)** | 0.172 (0.213) |
| CHK1 O14757 | 138 (126) | 4 | 3 (2) | 0.928 (0.992) | **0.329 (0.248)** | **0.068 (0.008)** | 0.088 (~0) |
| PIM1 P11309 | 153 (31) | 5 (4) | 3 (2) | 0.974 (0.903) | **0.115 (0.167)** | **0.011 (0.045)** | −0.008 (~0) |

Sup **Table ST2.** Agreement between intra-protein Louvain communities and independently assigned structural classes. Values include medium- and high-confidence annotations; values in parentheses use high-confidence annotations only. Site and structural-class counts refer to the corresponding annotated subsets.
