## supplementary text for "Network organization of protein binding-site landscapes across scales"

### 1 - Mapping of residue-contact patterns to six coarse structural classes

Residue-contact footprints mapped to canonical UniProt numbering were interpreted using kinase-specific structural landmarks, documented ligand-binding modes, and representative deposited structures. This curation distinguished contacts within the canonical ATP-binding cleft from extensions into adjacent subpockets and footprints associated with spatially distinct regulatory sites. Particular attention was given to the CK2α αD and α/β-interface pockets, the p38α C-terminal lipid pocket, the AURKA TPX2-binding Y-pocket, the CDK2 ANS-associated region, and the ERK2 αC/P-loop niche. For each binding site, we recorded the mapped residue footprint, mapping quality, structural assignment, confidence level, and residue-based rationale. These annotations constituted the curated reference set for interpreting intra-protein Louvain communities. Sites contacting the DFG region were designated back-pocket/DFG-associated, without inferring a DFG-out conformation from residue contacts alone.

### 2 - Description of the five connected-components of the distant-protein network mentioned in the text (Sup Fig SF7B)

Component 288 connected the class B G protein-coupled receptor (GPCR) GLP1R with four class A receptors: CB1, NTSR1, SUCNR1, and GPR174. These receptors share a seven-transmembrane architecture but differ in their evolutionary relationships and extracellular organization; GLP1R has a prominent extracellular domain (Kawai et al. 2020). Their small-molecule binding sites in the transmembrane regions nevertheless occupied nearby positions in embedding space. The retained edges connected ligand-contacting environments associated with the seven-transmembrane receptor architecture, suggesting shared local recognition features whose structural basis remains to be established. GLP1R was represented by several structurally characterized agonists, whereas the class A receptors were represented by chemically and pharmacologically diverse ligands. A single shared ligand identity therefore did not explain the component. The pattern is consistent with shared local recognition features and worth structural assessment.

Component 11 connected binding sites from enzymes in different metabolic pathways, including mammalian aldolase and phosphoglucomutase, bacterial phosphofructokinase, cofactor-independent phosphoglycerate mutase, and enzymes involved in organic-acid metabolism. The ligands included phosphorylated sugars, phospho-acids, and other polar metabolites or inhibitors. No single ligand accounted for the component. Recognition of polar or charged metabolites offers a possible explanation for the shared embedding features, but physicochemical convergence remains an interpretation requiring coordinate-level validation.

Component 53 connected phosphoacetylglucosamine mutase AGM1 from Candida albicans, ATIC enzymes involved in de novo purine biosynthesis, and ornithine decarboxylase from Trypanosoma brucei. These sites provided a related example involving chemically and pharmacologically diverse ligands. They involved phosphorylated N-acetylglucosamine metabolites in AGM1, synthetic inhibitors of the ATIC transformylase site, and the mechanism-based ornithine decarboxylase inhibitor DFMO/eflornithine. The proteins belong to different structural and functional families, yet their ligand-contacting environments were connected in the network. Shared recognition features for polar or charged compounds offer a possible explanation. This component also illustrates the limits of pharmacological extrapolation: eflornithine inhibition depends on the catalytic machinery of ornithine decarboxylase, and binding-site embedding similarity does not establish inhibition of the other proteins.

Component 175 contained 2-oxoglutarate-dependent oxygenases with different biological functions and InterPro assignments. All sites were associated with 2-oxoglutarate, providing a common biochemical basis for their similarity. This component is consistent with shared features of Fe(II)/2-oxoglutarate-dependent catalysis and provides a reference example of cosubstrate-associated network organization. Possible cross-reactivity of inhibitors targeting these sites requires independent pharmacological evaluation (Nakashima et al. 2021).

Component 72 contained enzymes recognizing adenosine-containing cofactors or metabolites, including proteins involved in CoA, cobalamin, and nucleotide-dependent chemistry. Recognition of related adenine–ribose–phosphate substructures provides a plausible biochemical explanation for these connections, illustrating how shared ligand features can contribute to binding-site similarity across different proteins.

These examples suggest that local binding-site similarity across globally distant proteins can have different biological origins. The connections provide hypotheses for shared recognition features and potential implications for off-target interactions or selectivity. Cofactor associated components, such as the 2-oxoglutarate oxygenases, have a readily identifiable shared chemical context. Components 11 and 53 and the cross-class GPCR component contain more heterogeneous ligand and protein sets, for which the molecular basis of similarity remains less directly resolved. As stated in the main text, the uncovered associations served for generating hypotheses about molecular recognition that should be challenged by structural and biochemical analyses in order to distinguish common chemical recognition, remote evolutionary relationships, physicochemical convergence, and pharmacologically transferable interactions.
